# An Approach to Measure Conductivity Gradients of Brain Tissue Transitions using Electrode Arrays

**DOI:** 10.64898/2026.07.10.737664

**Authors:** Markus Symmank, Moritz Gerber, Thomas Knösche, Erdem Güresir, Florian Wilhelmy, Konstantin Weise

## Abstract

Accurate determination of surgical margins is critical in tumor resection to ensure complete removal of tumor-infiltrated tissue while preserving healthy tissue. Pathological assessment provides reliable information but is time-consuming. This study investigates the feasibility of using impedance spectroscopy to detect tissue transitions at a macroscopic level. Two electrode arrays—one-dimensional and two-dimensional—were applied to ex vivo porcine brain tissue. Measurements were performed using both two- and four-electrode configurations, and data were corrected using the multiple-load compensation method. Results demonstrate that the one-dimensional array provides continuous conductivity profiles corresponding to tissue transitions, while the two-dimensional array showed less consistent results. These findings suggest that impedance spectroscopy is a promising tool for intraoperative margin detection, but further optimization of electrode geometry and measurement data processing is required.

## I. Introduction

Determining surgical margins during tumor resection presents a major challenge. The goal is to preserve as much healthy tissue as possible while completely removing tumor-infiltrated tissue [1]. Failure to remove all tumor tissue results in unsuccessful treatment [2]. Pathological assessment of resection margins during and after surgery provides the most reliable information about treatment success, but this process is time-consuming [3].

Since studies have shown that different tissue types exhibit distinct conductivities [4], [5], impedance spectroscopy could be used to identify conductivity gradients and thereby detect tissue transitions. This approach may allow detection of tumor infiltration zones and facilitate intraoperative determination of surgical margins.

The present study investigates the technical feasibility of this approach at a macroscopic level. Based on these results, the study highlights potential benefits and challenges, providing a foundation for further targeted development.

## II. Methods

### A. Measurement Setup

Measurements were performed using the *ISX-3* impedance analyzer (Sciospec, Bennewitz, Germany) in combination with a multiplexer, enabling rapid measurement with different electrode pairs. The multiplexer accepts boards with electrode arrays of various geometries. Figure 1(a) shows the hardware setup, and Figure 1(b) depicts a 3D CAD model of the board containing different electrode arrays. Two specific electrode arrays were used in this study.

**Fig. 1.**
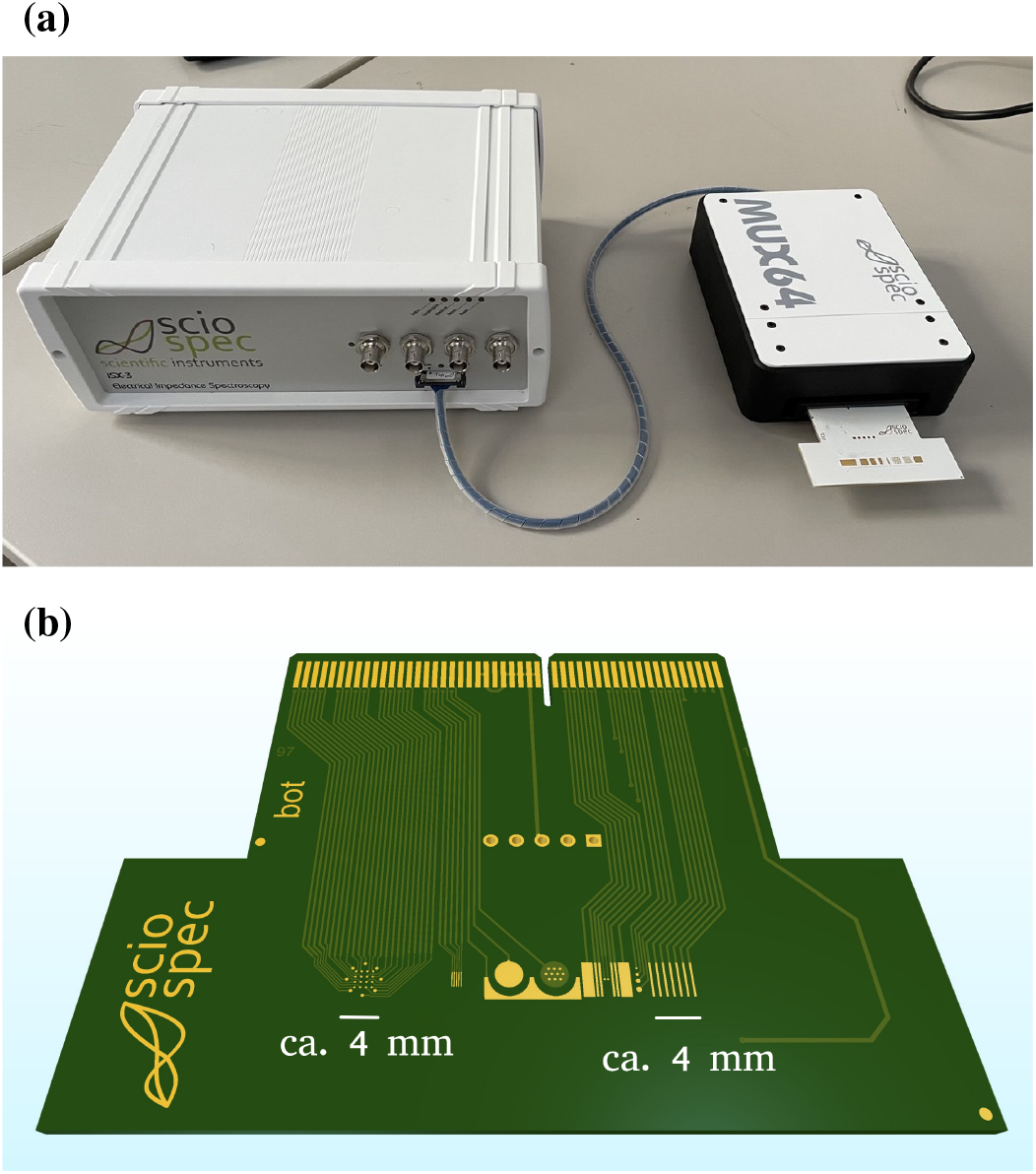
Measurement setup (a) with electrode arrays (b).

The first array consists of seven electrodes arranged in a comb-like, one-dimensional configuration (see Fig. 2(a)). The second array is a 4*×*4 equidistant matrix surrounded by a ring of eight electrodes distributed at 45° intervals (see Fig. 2(b)). Each array is used in both two- and four-electrode configurations. For the one-dimensional array in the four-electrode setup, the outer electrodes (16, 10) were used for current injection, while the voltage was measured sequentially between adjacent inner electrode pairs (15,14), (14,13), (13,12), and (12,11), from which the corresponding impedances were calculated. Two-electrode measurements were conducted using the same sensing electrode pairs for comparability. In this case the excitation electrodes are omitted.

**Fig. 2.**
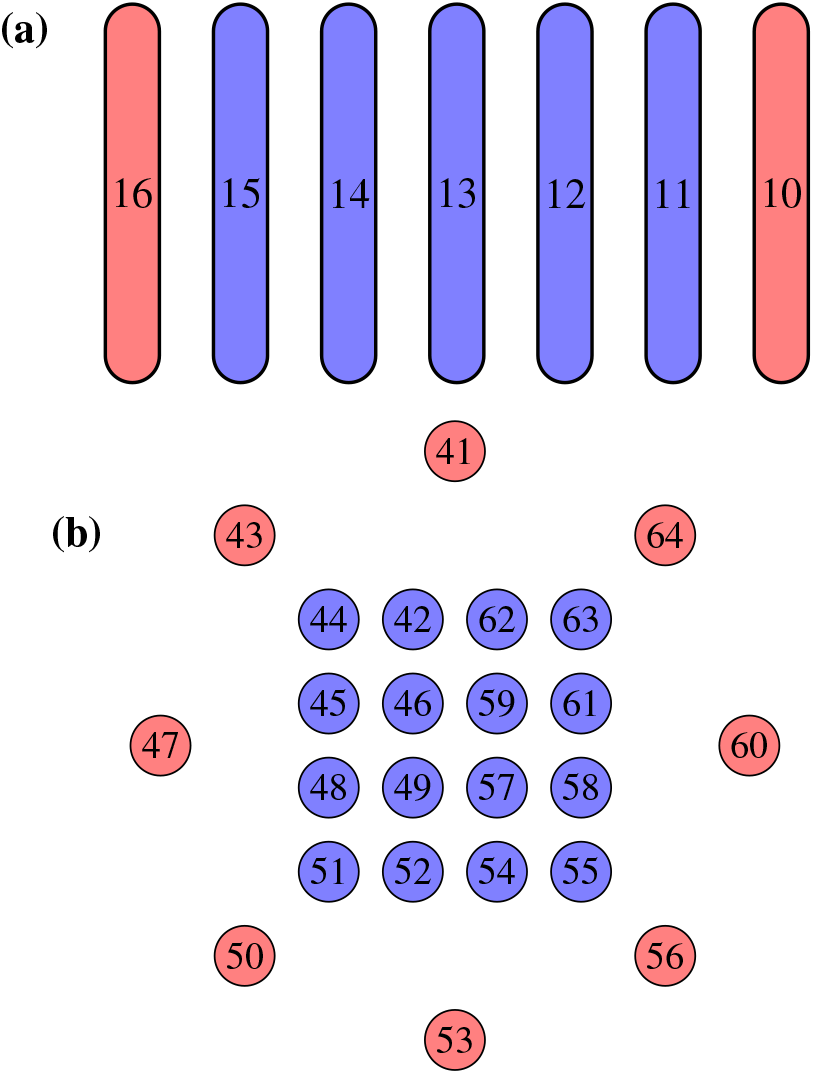
Schematic of used two-dimensional electrode arrangement.

For the two-dimensional array, four-electrode measurements were performed using current injection through pairs of opposite electrodes in the outer ring: (41,53), (64,50), (60,47), and (56,43). Impedance was measured between vertically adjacent matrix electrodes in a row-wise sequence. This resulted in twelve measurement positions. However, four-electrode measurements in this geometry produced unreliable data, so the focus was placed on two-electrode measurements, following the same row-wise order while the outer ring remained unused.

### B. Calibration

The calibration and compensation procedure follows the method described in [6], adapted for electrode arrays. The steps of the calibration, which are also illustrated in Fig. 3, consist of the following:

1. **Measurement of calibration solutions:** KCL-Solutions with varying dilutions and conductivities were applied as droplets on the electrode arrays. Impedance spectra were recorded with 30 repetitions per measurement. KCl solutions were selected as calibration electrolytes due to their well-defined and stable conductivity, predominantly ohmic behavior over the investigated frequency range, low electrode polarization effects, and widespread use in bioimpedance calibration protocols. This choice ensures reproducibility and comparability with established literature and commercially available conductivity standards.
2. **Reference measurement:** Conductivities of the calibration solutions were measured using a conductivity meter (ScioSpec, TetraCon 325 probe). These values served as references for further calibration.
3. **Selection of reference frequencies:** Reference frequencies were chosen where the impedance is approxiin magnitude was selected. Unlike single-point measurements, reference frequencies for electrode arrays vary depending on electrode pair, measurement method, and solution.
4. **Multiple-load compensation:** Measured impedances were compensated using the multiple-load compensation method described in [6]. The calibration solutions used for compensation included 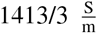, 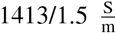, and 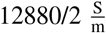 These solutions were selected such that their impedances span the range expected during brain tissue measurements. Although the compensation method is based on a two-port model, previous studies [6] indicate that it improves data from four-electrode measurements in a manner comparable to its effect on two-electrode measurements.
5. **Determination of the geometry factor:** Compensated impedances of the solutions were converted to admittances at the reference frequencies. The real part of the admittance (conductance) for each solution was plotted against its reference conductivity, forming a scatter plot. Linear regression provided the slope, which corresponds to the inverted geometry factor 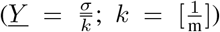. This factor allows conversion of measured conductance to specific tissue conductivity.

**Fig. 3.**
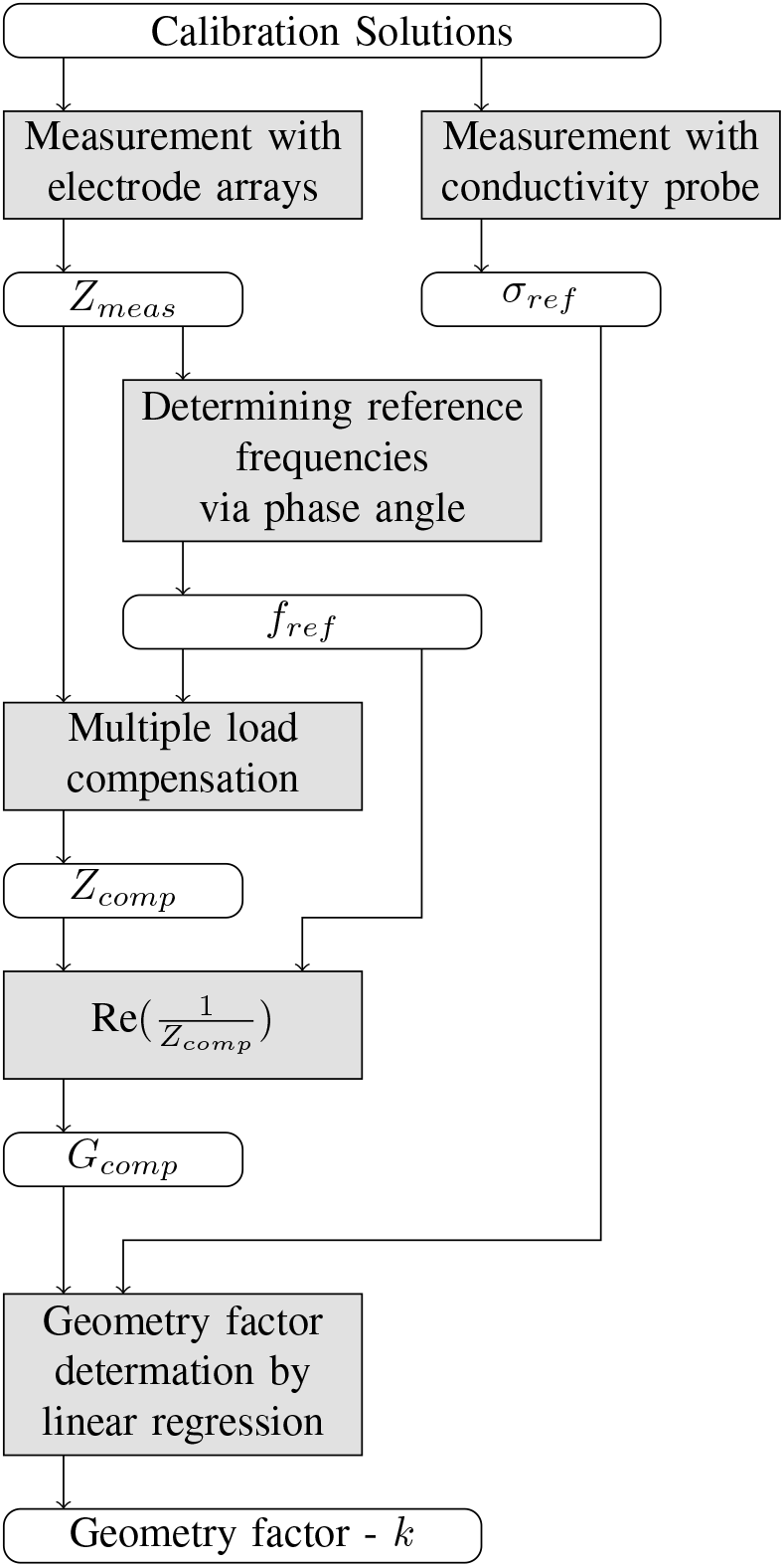
Calibration procedure of measuring chain using calibration solutions.

### C. Porcine Brain Tissue Measurement Procedure

For the impedance measurements, the impedance analyzer was connected to the multiplexer, together with the PCB carrying the electrode arrays. The measurement specimen consisted of an ex vivo brain sample obtained from a domesticated pig. A tissue sample was excised from the transition zone between grey and white matter and positioned on the electrode array. The experimental setup is shown in Fig. 4.

**Fig. 4.**
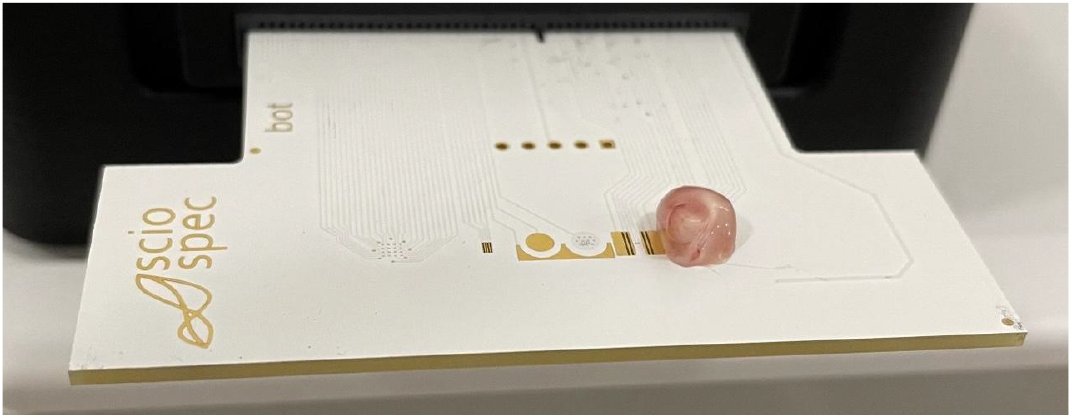
Tissue placement of grey to white matter transition on electrode array.

Using the previously acquired calibration data, the raw measurements of the tissue samples were first compensated and subsequently converted into specific conductivities. The individual processing steps are summarized schematically in Fig. 5. The measurement procedure was performed twice for each electrode array and configuration. The orientation of the transition from grey to white matter in the second run is exactly opposite to the orientation in the first run. This approach was used to verify that the resulting conductivity gradients were not caused by systematic measurement bias.

**Fig. 5.**
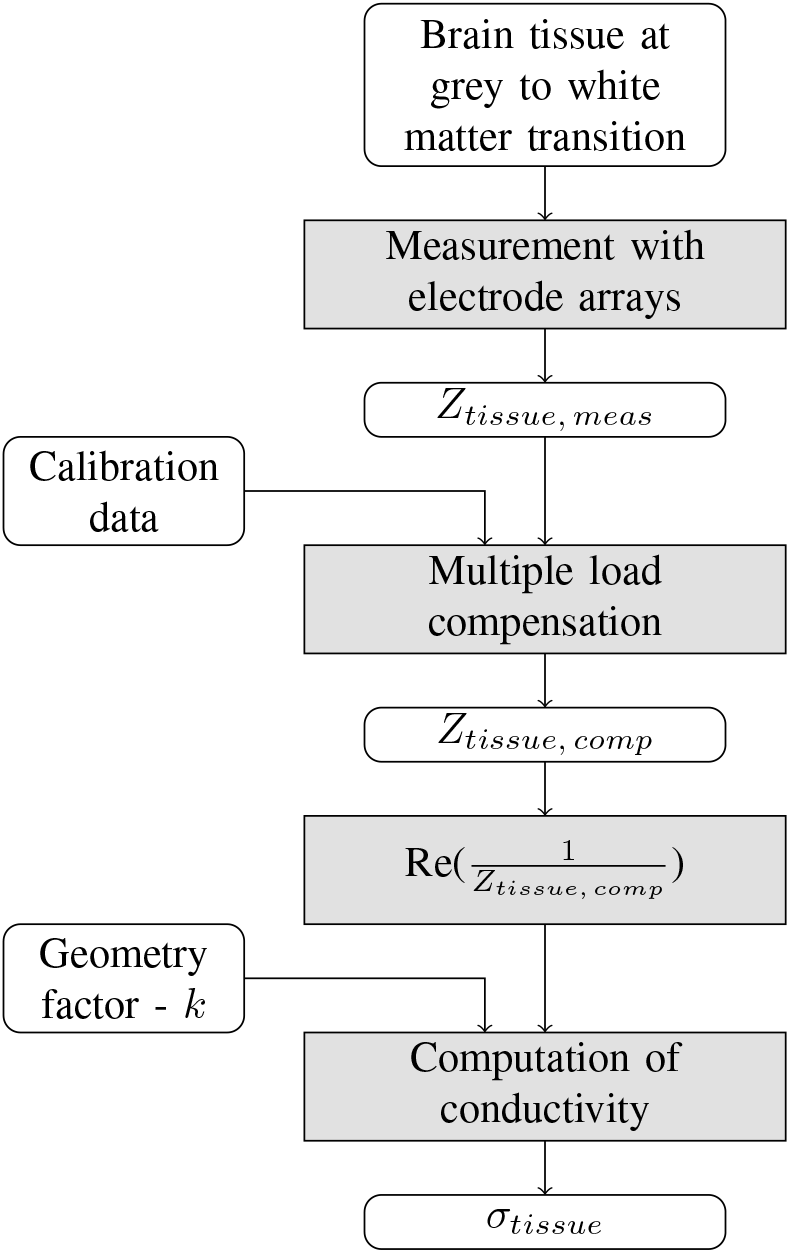
Brain tissue measurement and data processing procedure.

## III. Results

### A. Precision in Calibration Solutions Measurement

For each calibration solution, 30 repeated measurements were performed for every electrode array and for every electrode configuration. Representative results are presented in terms of the measured mean values and standard deviations. Fig. 6 shows the results obtained for calibration solution 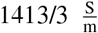 for the one-dimensional array (a) excitation electrodes: (16, 10); sensing electrodes: (15, 14) and the two-dimensional array (b) excitation electrodes: (41, 53); sensing electrodes: (45, 48). The one-dimensional array exhibits high repeatability in both the two-electrode and four-electrode configurations (Fig. 6(a)), as indicated by the low standard deviation across the 30 repetitions. The same behavior is observed for the two-dimensional array when operated in the two-electrode configuration. In contrast, operation of the two-dimensional array in the four-electrode configuration (Fig. 6(b)) yields a substantially higher standard deviation relative to the mean. This behavior serves as an initial indication that measurements obtained with the two-dimensional array in four-electrode mode may be influenced by yet unidentified factors.

**Fig. 6.**
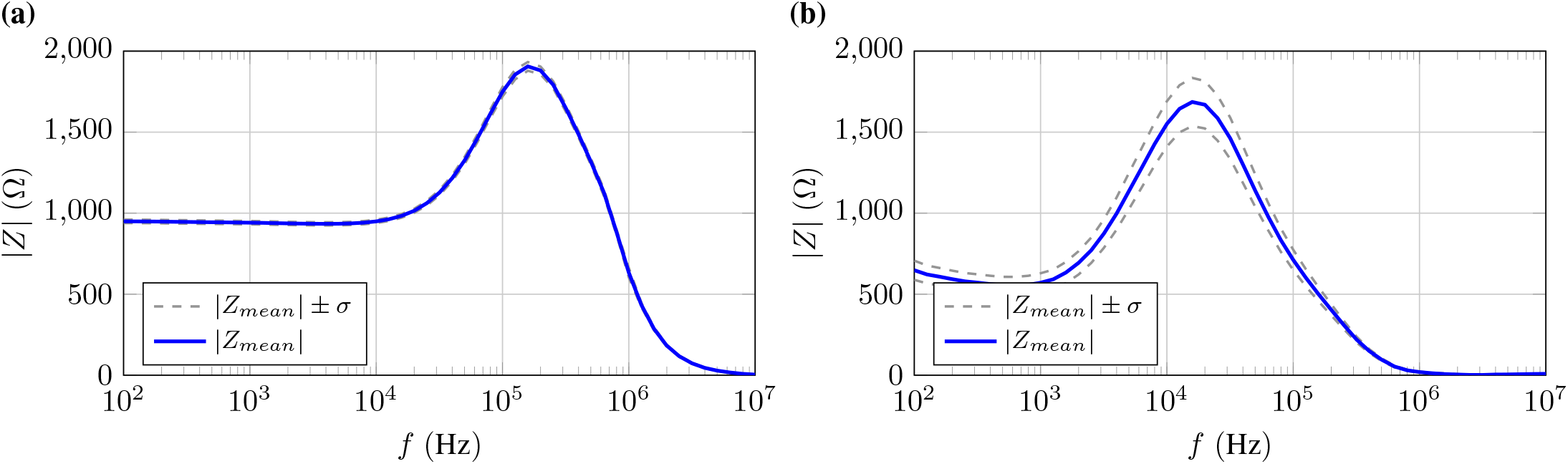
Mean and standard deviation of the absolute value of the impedance obtained from 30 repeated measurements of calibration solution 1413/3 using four-electrode configuration with (a) one-dimensional electrode array (b) two-dimensional electrode array.

### B. Compensation of Calibration Solutions

Validation of the multiple-load compensation [6] was performed by comparing the measured and compensated results of all calibration solutions. The corresponding results are shown in Fig. 7. The three calibration solutions used as reference impedances for the compensation appear as horizontal lines in the spectrum. Solutions whose impedances lie between those of the reference solutions impedances also exhibit substantial smoothing after compensation. Fig. 7 illustrates the impedances measured with the one-dimensional electrode array in the four-electrode configuration (excitation electrodes: (16, 10); sensing electrodes: (15, 14)) both (a) before and (b) after compensation. The same comparison is shown for the two-dimensional array in the two-electrode configuration (electrodes: (45, 48)) in (c) before and (d) after compensation.

**Fig. 7.**
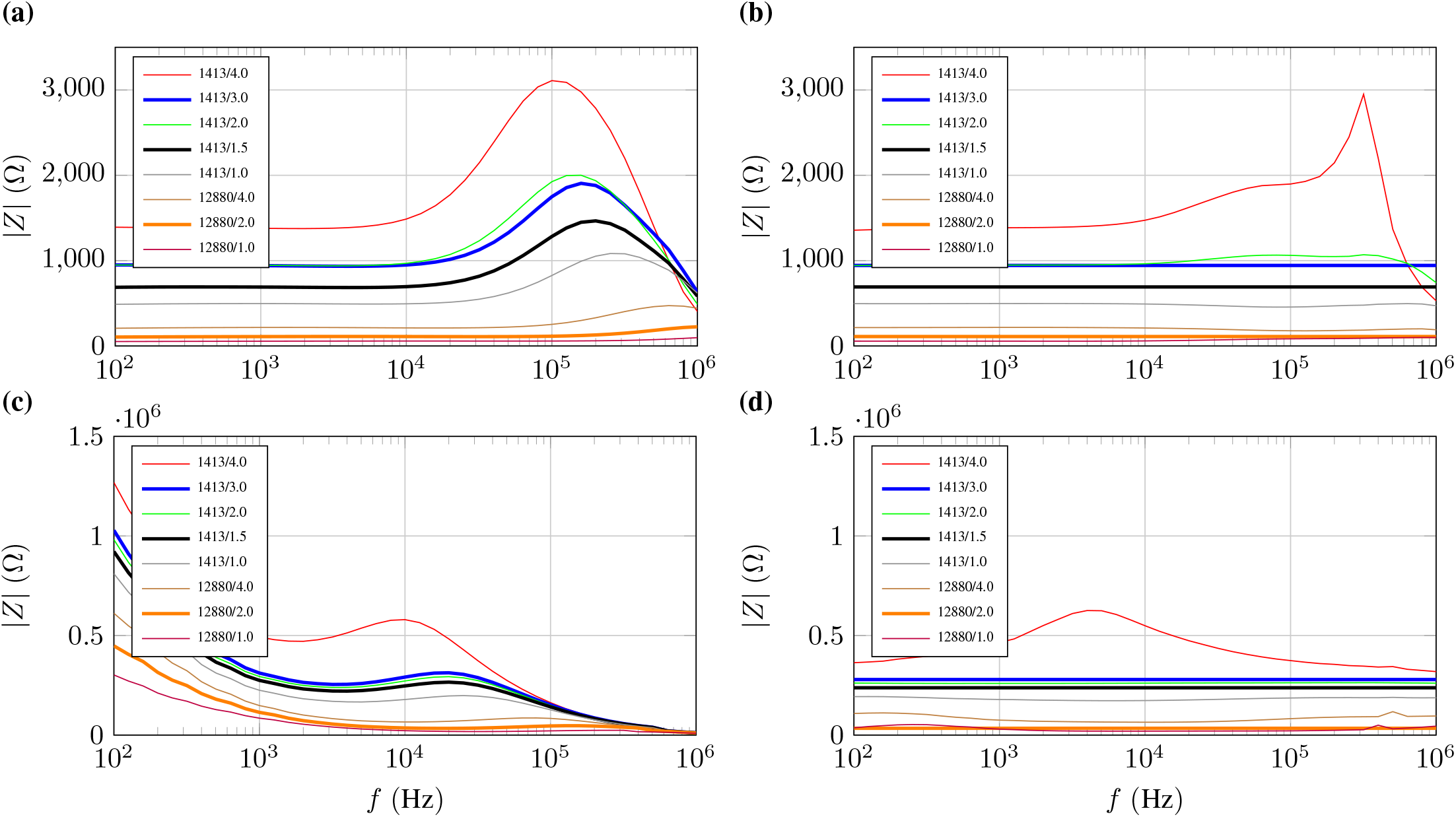
Magnitude impedance data of calibration solutions for the one-dimensional array in the four-electrode configuration (a) before and (b) after compensation, and for the two-dimensional array in the two-electrode configuration (c) before and (d) after compensation.

### C. Geometry Factor Determination

The geometry factors were determined individually for each electrode configuration. Figure 8 shows representative regression lines whose slopes correspond to the inverse of the geometry factor, illustrated for (a) the one-dimensional array in the four-electrode configuration (excitation electrodes: (16, 10); sensing electrodes: (15, 14)) and (b) the two-dimensional array in the two-electrode configuration (electrodes: (45, 48)). The calibration data obtained with the two-dimensional array in the four-electrode configuration are inconsistent and linear regression failed for nearly all electrode configurations. Table I summarizes the minimum and maximum *R*^2^ and *p* values obtained for both array types under two-electrode and four-electrode measurement conditions.

**Fig. 8.**
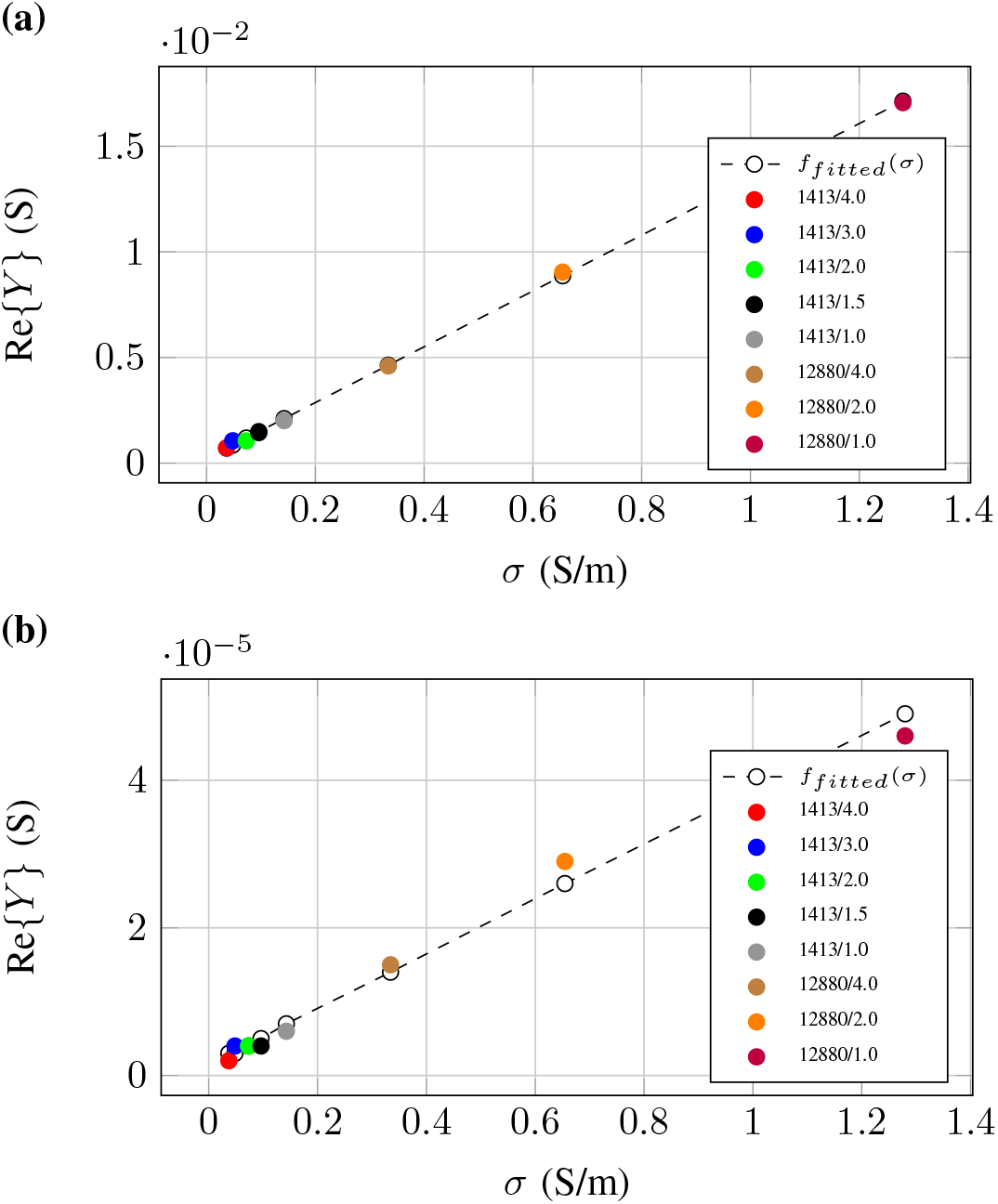
Geometry factor determination by regression line for (a) the one-dimensional array in the four-electrode configuration and (b) the two-dimensional array in the two-electrode configuration.

**TABLE I.** Geometry Factor Regression Results.

| Geometry / Method | $R_{min}^2$ | $R_{max}^2$ | $p_{min}$ | $p_{max}$ |
| --- | --- | --- | --- | --- |
| one-dim. / two-electrodes | 0.996 | 0.997 | < 0.001 | < 0.001 |
| one-dim. / four-electrodes | 0.999 | 0.999 | < 0.001 | < 0.001 |
| two-dim. / two-electrodes | 0.904 | 0.997 | < 0.001 | < 0.001 |
| two-dim. / four-electrodes | 0.010 | 0.981 | < 0.001 | 0.817 |

### D. Grey to White Matter Transition Measurement

The data obtained from the tissue measurements are visualized in Fig. 9 for the one-dimensional array. A clear conductivity gradient is observed, and the orientation of the grey-to-white matter transition is distinctly reflected in the measurement data. This confirms that the results are not caused by systematic errors in the instrumentation or measurement procedure. Instead, the observed variations represent genuine conductivity differences arising from the respective tissue types.

**Fig. 9.**
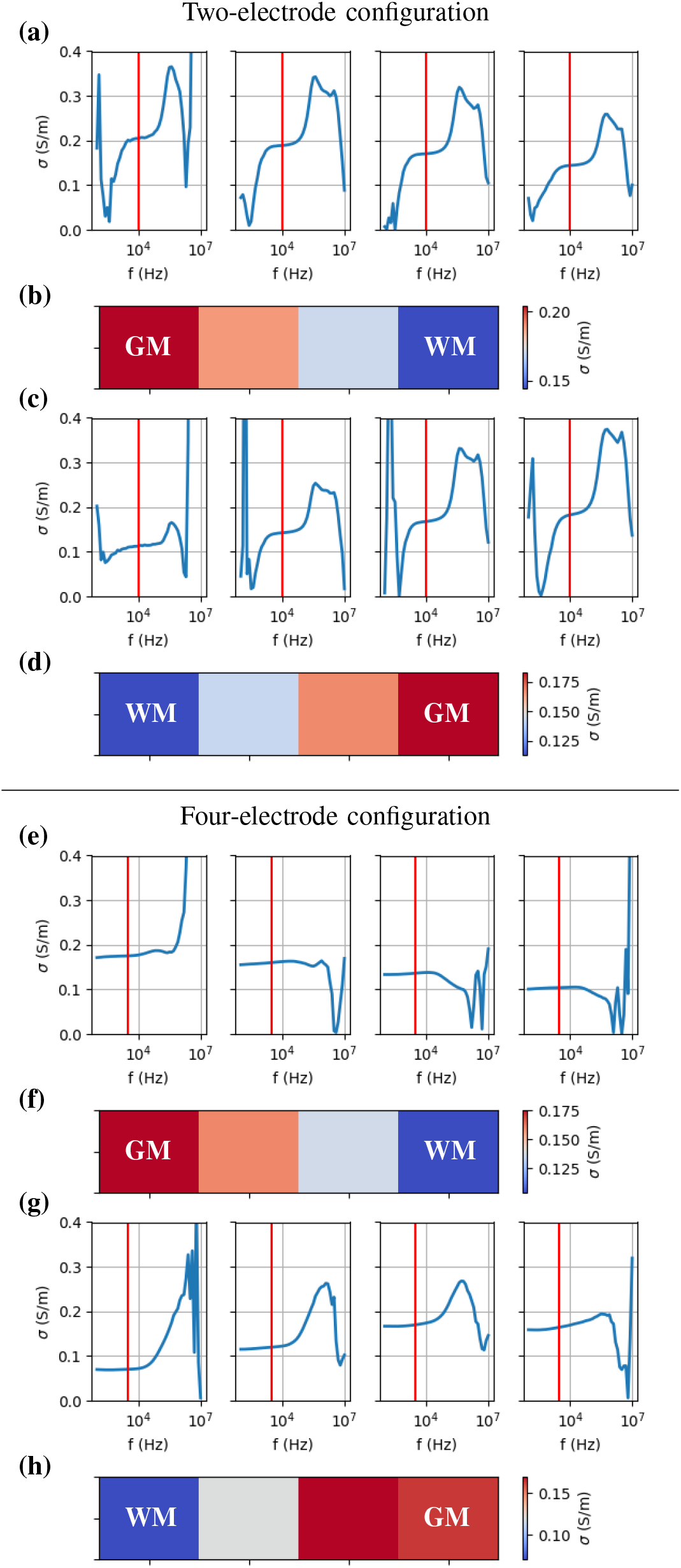
Measurement results of brain tissue transition with one-dimensional electrode array: (a, b) grey to white matter in two-electrode configuration; (c, d) white to grey matter in two-electrode configuration; (e, f) grey to white matter in four-electrode configuration; (g, h) white to grey matter in four-electrode configuration; (b, d) conductivity gradient at 10 kHz; (f, h) conductivity gradient at 3 kHz.

The situation differs for measurements performed with the two-dimensional array. In neither the two-electrode nor the four-electrode configuration could the tissue transition be identified in the resulting conductivity values. The results obtained for the two-electrode measurements are shown in Fig. 10.

**Fig. 10.**
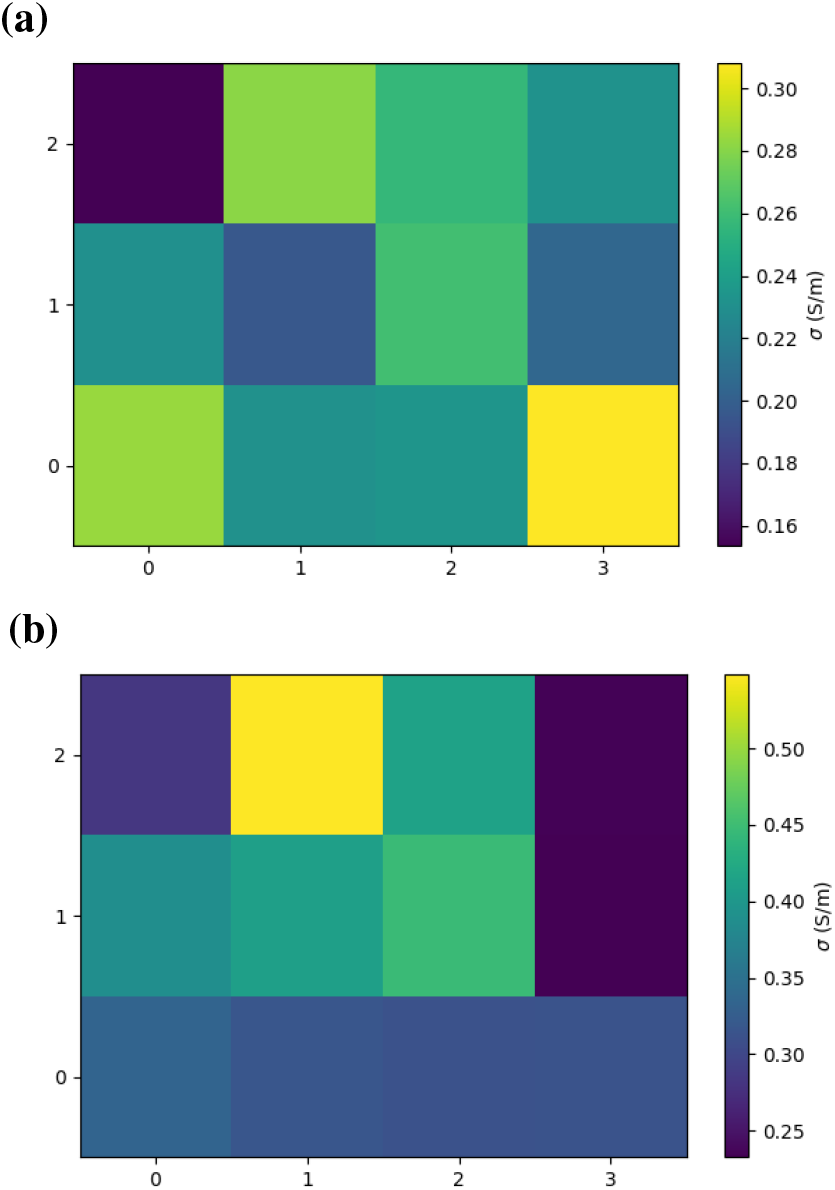
Measurement results of brain tissue transition with two-dimensional electrode array in two-electrode configuration: (a) grey to white matter; (b) white to grey matter.

### IV. Discussion AND Future Work

The results indicate that the approach pursued here is indeed capable of resolving a macroscopic conductivity gradient. At the same time, it becomes evident that the measurement procedure requires further investigation. The findings obtained with the two-dimensional array suggest that this geometry may introduce substantial electric-field inhomogeneities, which have a decisive impact on data quality. In particular, the outer ring of larger electrodes surrounding the inner matrix may lead to scattering effects that are difficult to predict. Additional influencing factors including the parallel conductor traces on the PCB and the uncertain impedance characteristics of the multiplexer. The strength of these effects is currently unclear.

A direct comparison between two-dimensional measurements and consecutive one-dimensional scans was not performed in this study. While such a comparison appears conceptually appealing, it is not trivial due to fundamentally different current distributions and electric field inhomogeneities introduced by the dense electrode geometry of the two-dimensional array. In particular, for four-electrode configurations, transversal impedance effects and uncontrolled coupling paths may significantly influence the measured signals. A systematic investigation of this comparison is therefore considered an important subject of future work.

Future work should aim to minimize the identified sources of uncertainty and to validate the measurements histologically. A promising direction may be the development of a mesoscopic electrode array fabricated directly on a glass substrate, enabling measurements on histological sections under an epi-fluorescence or bright-field microscope. Such a smaller-scale geometry may reduce scattering effects. Another potential improvement includes driving the excitation through multiple adjacent point electrodes to promote a more homogeneous electric field.

Furthermore, future electrode arrays should feature a higher-resolution electrode grid in order to identify additional measurement configurations and potentially enable the determination of not only individual tissue transitions but also cortical layers at the mesoscopic level [7]. The results suggest that the one-dimensional array is particularly suitable for this application.

### V. Conclusion

This study demonstrates the technical feasibility of using impedance spectroscopy for detecting tissue transitions in ex vivo porcine brain tissue. One-dimensional electrode arrays provided reliable and continuous conductivity profiles, while two-dimensional arrays require further optimization. These findings support the potential of impedance-based methods for intraoperative margin detection. Further development, including array optimization, is necessary to translate this approach to clinical practice.

## VI. Acknowledgement

The project was greatly supported by Sciospec Scientific Instruments GmbH (04828 Bennewitz, Germany) providing rental equipment.

The authors used an AI-based language model (ChatGPT, OpenAI) to assist in language refinement.

## Notes

### Competing Interest Statement

The authors have declared no competing interest.

